## Supplemental Figure 1 for "A conserved glutathione binding site in poliovirus is a target for antivirals and vaccine stabilisation"

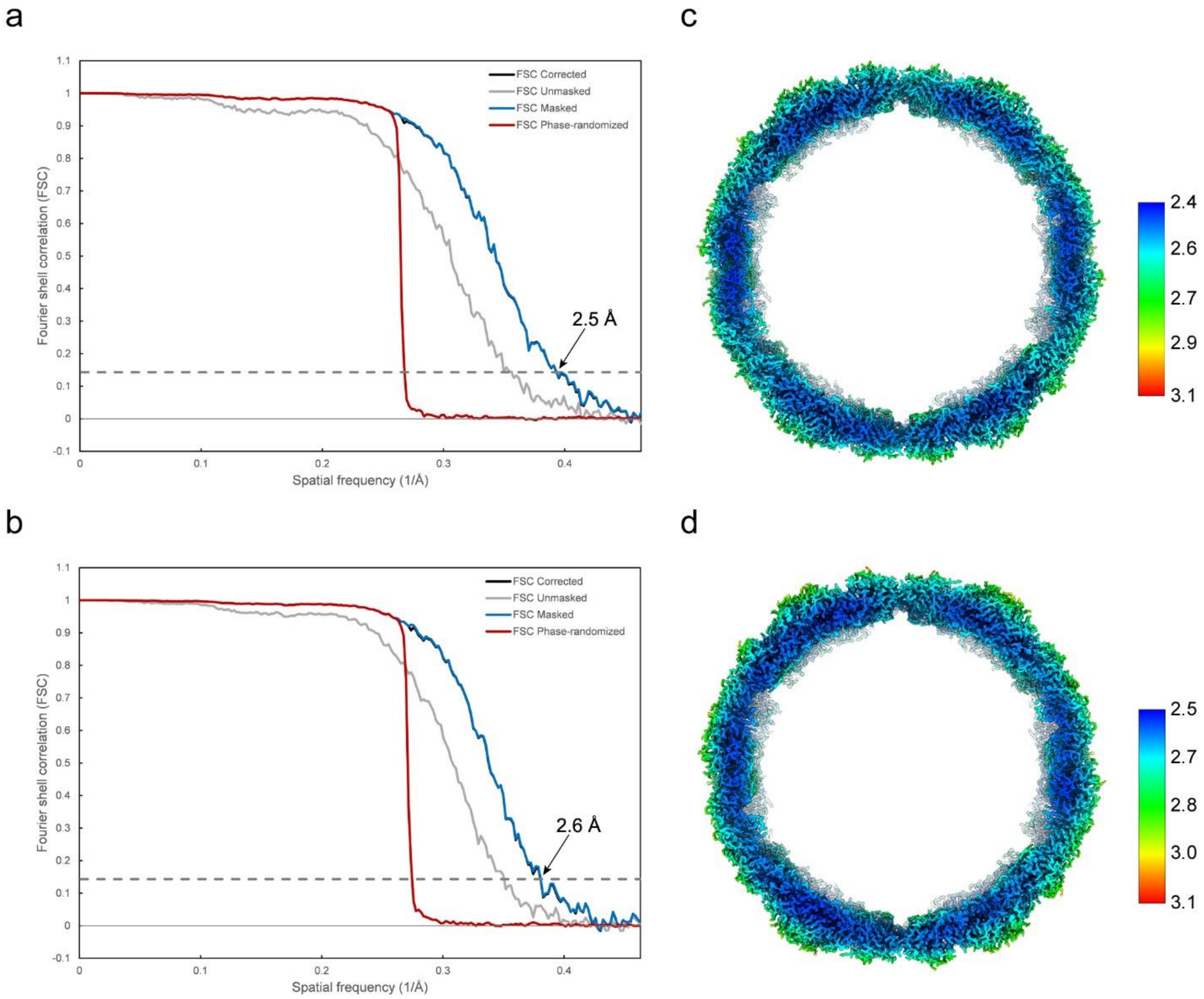


**Supplementary Figure 1. Resolution of the PV3-SC8$^{GPP3+GSH}$ and PV3-SC8$^{pleconaril+GSH}$ reconstructions.** **a,b** Fourier shell correlation (FSC) calculated between two independent half sets of data as a function of spatial frequency is plotted for the PV3-SC8$^{GPP3+GSH}$ and PV3-SC8$^{pleconaril+GSH}$ reconstructions respectively. FSC is plotted for the original unmasked half-maps (grey) and masked half-maps that had density corresponding to solvent removed (blue). FSC is also shown for phase-randomized half-maps (red) used to compensate for possible effects of the masking procedure before calculating the final corrected FSC (black). Good agreement between the masked and corrected curves indicated no adverse effects from the masking. The resolution at which the corrected curve drops below the FSC=0.143 threshold (grey dashed line) is indicated with an arrow. **c,d** Local resolution analysis of the final cryo-EM electron potential maps for PV3-SC8$^{GPP3+GSH}$ and PV3-SC8$^{pleconaril+GSH}$ reconstructions, respectively as assessed by RELION local resolution estimation. A central slice through the VLPs is viewed along the 2-fold axis and the distribution of local resolution is shown coloured from blue to red according to the colour key shown.
