## Supplemental Figure 2 for "A conserved glutathione binding site in poliovirus is a target for antivirals and vaccine stabilisation"

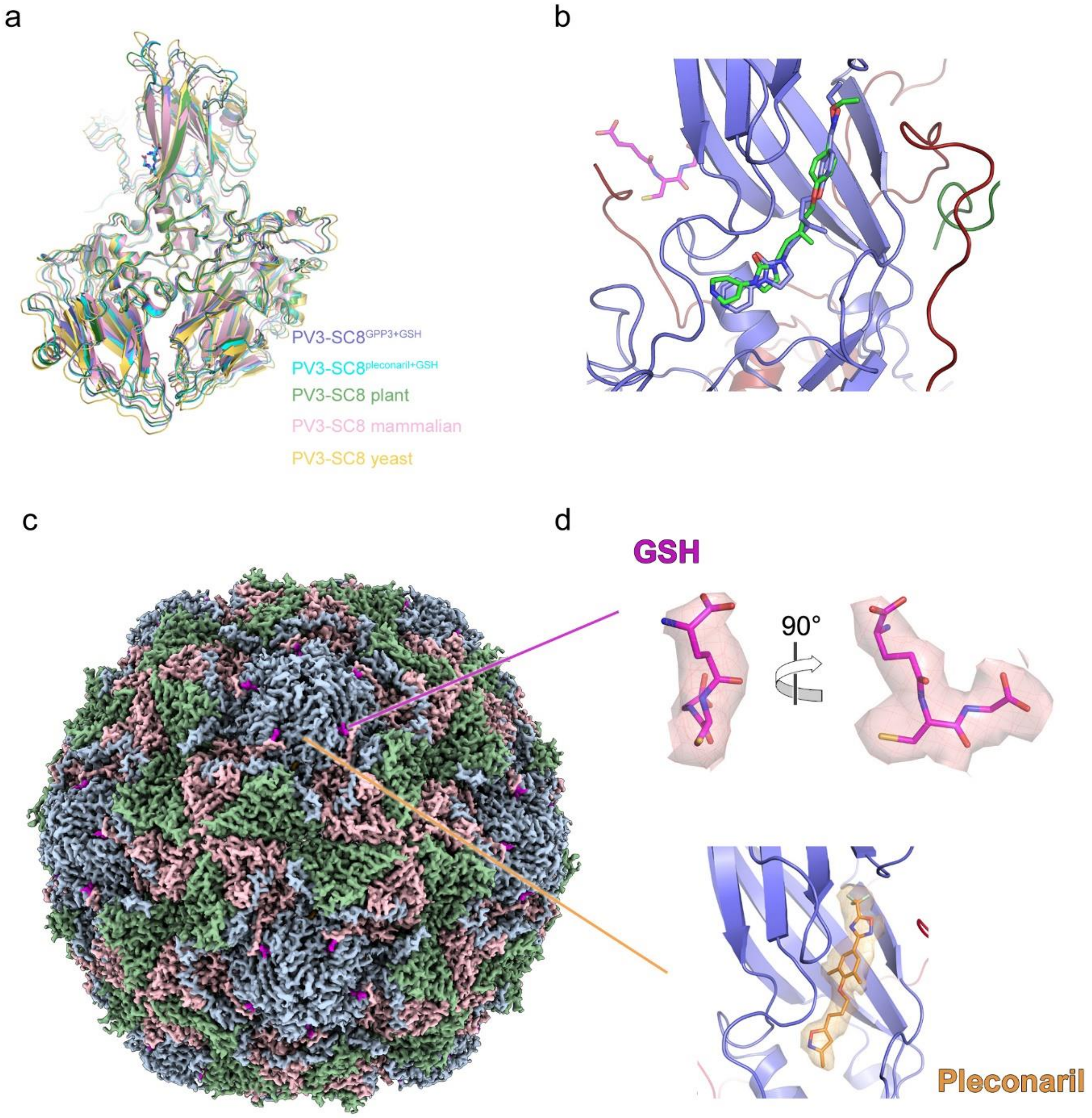


**Supplementary Figure 2. Similarity of PV3-SC8 structures, GPP3 binding mode and PV3-SC8$^{\text{pleconaril+GSH}}$ reconstruction.** **a** Structure superposition of the capsid protomers of the PV3-SC8$^{\text{GPP3+GSH}}$ complex (blue) with PV3-SC8$^{\text{pleconaril+GSH}}$ (cyan), apo PV3-SC8 (yellow), PV3-SC8 from plant cell expression (green) {Marsian, 2017 #134} and PV3-SC8 from mammalian cell expression (pink) {Bahar, 2021 #5}. **b** VP1 pocket of the PV3-SC8$^{\text{GPP3+GSH}}$ comparing binding mode of bound GPP3 (blue sticks) with GPP3 bound in plant cell produced PV3-SC8 (green sticks). **c** Three-dimensional reconstruction of PV3-SC8 VLP after incubation with a molar excess of GSH and pleconaril. The VLP is viewed along the icosahedral twofold symmetry axis with the VP1, VP0 and VP3 subunits of the capsid protomer coloured light blue, light green and light red respectively. **d** Expanded views of bound GSH and pleconaril fitted into the cryo-EM electron potential map. For GSH the electron potential map is shown at 1 σ and for pleconaril at 1.5 σ. All maps are rendered at a radius of 2 Å around atoms.
