## Supplemental Figure 3 for "A conserved glutathione binding site in poliovirus is a target for antivirals and vaccine stabilisation"

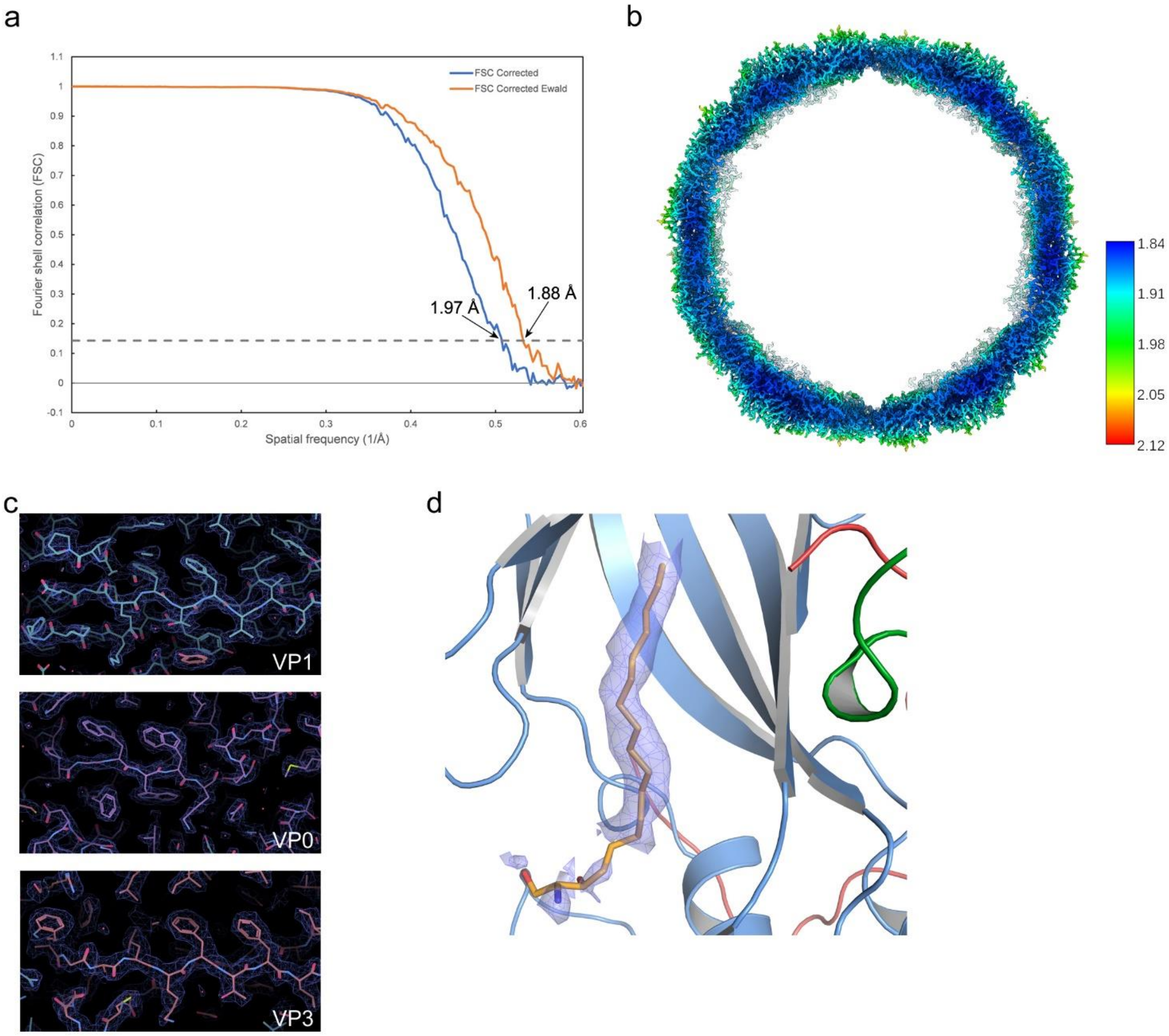


**Supplementary Figure 3. Resolution of the wt PV2-CP17 reconstruction before and after Ewald sphere correction and density features. a** Fourier shell correlation (FSC) calculated between two independent half sets of data as a function of spatial frequency is plotted for the wt PV2-CP17 reconstruction. The final corrected FSC is plotted for reconstructions before (blue) and after (orange) Ewald sphere correction. The resolution at which the corrected curve drops below the FSC=0.143 threshold (grey dashed line) is indicated with an arrow. **b** Local resolution analysis of the final cryo-EM electron potential map for the wt PV2-CP17 reconstruction as assessed by RELION local resolution estimation.  A central slice through the reconstruction is viewed along the icosahedral 2-fold axis and the distribution of local resolution is shown coloured from blue to red according to the scale bar shown.  c Representative snapshots of cryo-EM density map for the VP1, VP0 and VP3 subunits of the cryo-EM electron potential map for the wt PV2-CP17 reconstruction after Ewald sphere correction and sharpening with a *B*-factor of -15 Å$^2$.  **d** View of sphingosine modelled into the VP1 pocket of the wt PV2-CP17 reconstruction.  The cryo-EM electron potential map is shown at 1 $\sigma$ for the modelled sphingosine and rendered at 2 Å around atoms.  The cryo-EM density around the hydrophilic head domain of the sphingosine was disordered.
