## Supplemental Table 1 for "A conserved glutathione binding site in poliovirus is a target for antivirals and vaccine stabilisation"

**Supplementary Table 1.** Cryo-EM data collection and image processing

| | PV3-SC8$^{GPP3+GSH}$ | PV3-SC8$^{pleconaril+GSH}$ | wt PV2-CP17 |
|---|---|---|---|
| **Data Collection** | | | |
| Voltage (kV) | 300 | 300 | 300 |
| Magnification (×) | 129629 | 129629 | 60314 |
| Defocus range (μm) | -2.9 to -0.8 | -2.9 to -0.8 | -2.3 to -0.8 |
| Dose rate (e–/pixel/s) | 0.59 | 35.79 | 14.02 |
| Frames | 60 | 25 | 50 |
| Frame length (s) | 1.163 | 0.046 | 0.034 |
| Total electron dose ($e^{-}$/Å$^2$) | 34.89 | 35.29 | 34.68 |
| Micrographs | 2503 | 4706 | 4750 |
| **Data processing** | | | |
| Pixel size (Å) | 1.08 | 1.08 | 0.829 |
| Initial particles (no.) | 19622 | 34523 | 157002 |
| Final particles (no.) | 5364 | 15275 | 51518 |
| Box size (pixels) | 450 | 450 | 450 |
| Symmetry | I1 | I1 | I1 |
| Accuracy of rotations (°) | 0.1325 | 0.1845 | 0.1085 |
| Accuracy of translations (Å) | 0.2160 | 0.3175 | 0.1658 |
| Resolution (Å) | 2.54 | 2.64 | 1.88 |
| Map sharpening *B*-factor (Å$^2$) | -52.3 | -72.2 | -15.0 |
