## Supplemental Table 2 for "A conserved glutathione binding site in poliovirus is a target for antivirals and vaccine stabilisation"

**Supplementary Table 2.** Structure refinement and validation for the capsid protein (VP0, VP1, VP3)

| | PV3-SC8$^{GPP3+GSH}$ | PV3-SC8$^{pleconaril+GSH}$ | wt PV2-CP17 |
|---|---|---|---|
| **Model composition** | | | |
| Non-hydrogen atoms | 5863 | 5857 | 6565 |
| Protein residues | 736 | 736 | 802 |
| Ligands | GPP3:1 | Pleconaril:1 | Sphingosine:1 |
| | GSH:1 | GSH:1 | CP17:1 |
| Waters | | | 262 |
| **Refinement** | | | |
| Resolution (Å) | 2.54 | 2.64 | 1.88 |
| Map CC[a] (Mask) | 0.85 | 0.87 | 0.90 |
| Map CC[a] (Volume) | 0.83 | 0.85 | 0.88 |
| **RMS deviations** | | | |
| Bond lengths (Å) | 0.002 | 0.003 | 0.003 |
| Bond angles (°) | 0.470 | 0.455 | 0.549 |
| **Mean B-factor (Å$^2$)** | | | |
| Protein | 20.36 | 23.74 | 24.16 |
| Ligand | 19.37 | 23.69 | 24.60 |
| Water | | | 23.26 |
| **Validation** | | | |
| Molprobity[b] score (percentile) | 1.02 ($100^{th}$) | 0.91 ($100^{th}$) | 1.11 ($100^{th}$) |
| Clashscore[b], all atoms (percentile) | 2.42 ($99^{th}$) | 1.65 ($99^{th}$) | 2.42 ($99^{th}$) |
| Ramachandran favoured (%) | 98.20 | 98.20 | 97.60 |
| Ramachandran allowed (%) | 1.80 | 1.80 | 2.40 |
| Ramachandran outliers (%) | 0.00 | 0.00 | 0.00 |
| Rotamer favoured (outliers) (%) | 98.29 (0.16) | 98.13 (0.31) | 98.69 (0.15) |
| Cβ deviations >0.25 Å (%) | 0.00 | 0.00 | 0.00 |
| CaBLAM outliers (%) | 1.12 | 1.26 | 1.28 |
| CA Geometry outliers (%) | 0.56 | 0.28 | 0.26 |
| **EMRinger[c] score** | 4.99 | 4.90 | 7.84 |

[a] Map CC is given for the full particle reconstruction.

[b] Chen *et al.* (2010) Acta Crystallographica D66:12-21.

[c] Barad *et al.* (2015) Nature Methods 12:943–946.
